# Individual-specific effects of transcranial electrical stimulation on 40-Hz auditory steady-state responses

**DOI:** 10.1101/2025.01.29.635512

**Authors:** Aurimas Mockevičius, Xiaojuan Wang, Jovana Bjekić, Marko Živanović, Saša R. Filipović, Inga Griškova-Bulanova

## Abstract

Transcranial electrical stimulation (tES) has shown promise for modulating brain function and related behavioral performance, but evidence has been mixed thus far. The possibility of tracking brain activity changes following tES via neurophysiological markers would benefit a better understanding of tES effects and the future development of tES protocols. One promising marker is the auditory steady-state response (ASSR), an externally controlled oscillatory brain activity, typically at 40 Hz, evoked by a periodic auditory stimulus. This study examined the offline effects of different types of tES on 40-Hz ASSR. Participants underwent four conditions of tES, which were applied over the left posterior parietal cortex: transcranial direct current (tDCS), transcranial alternating current (tACS), oscillatory transcranial direct current (otDCS) and sham stimulation. Individually determined theta frequency was delivered in the tACS and otDCS protocols. Following the tES application, electroencephalogram (EEG) was recorded during 40-Hz auditory click stimulation. Mixed-effects modeling revealed no significant group-level differences in phase locking or evoked amplitude between stimulation conditions. However, both baseline (sham) ASSR and the change in ASSR following tES had a substantial interindividual variability. Exploratory analysis showed that individuals with lower baseline ASSR had increased synchronization following tES. Furthermore, the increase in ASSR synchronization was linked to higher memory gain; however, the relationship was observed only in otDCS condition. The findings encourage future research focusing on individual factors that may contribute to tES outcomes.

## I. INTRODUCTION

In recent years, advancements have been made in the application of noninvasive brain stimulation in the treatment of neuropsychiatric disorders [1]. In particular, transcranial electrical stimulation (tES) has gained considerable attention due to its favorable safety profile and ease of use [2]. TES delivers low intensity electrical current via scalp electrodes to modulate brain activity. Various types of tES, including transcranial direct current stimulation (tDCS), transcranial alternating current stimulation (tACS) and oscillatory transcranial direct current stimulation (otDCS) are thought to exert different modulatory effects on cortical excitability. In particular, tDCS modulates the resting membrane potential by applying a constant current between two opposing electrodes, thereby increasing cortical excitability [3]. TACS entrains neural oscillations to the stimulation frequency by rhythmically altering the polarity of the electrodes [4]. Meanwhile, otDCS combines features of tDCS and tACS, with the current oscillating at a specific frequency while maintaining the polarity of electrodes. This approach is thought to induce changes in both cortical excitability and oscillatory brain activity [5].

However, all tES modalities when they are applied using a conventional two-electrode setup are characterized by low spatial focality [6], [7], [8], [9]. Specifically, when two relatively large electrodes (e.g., 25 cm^2^) are placed on the scalp, the resulting electric field affects not only the targeted brain region but also adjacent areas due to the spatial spread of current. Moreover, the induced electric field is not confined to the cortical surface, and it extends to deeper brain structures, particularly in configurations involving extracephalic and cross-hemispheric electrode placement [10]. Finally, the physiological effects of tES are not restricted to the area beneath the electrodes, as stimulation effects are spread across functionally connected brain regions via network-propagation mechanisms [11]. This raises important questions about unintended physiological effects and highlights the need to investigate outcomes beyond the primary cognitive or behavioral outcomes. Moreover, it has also been proposed that tES effects can be explained by changes in the E/I balance [12], [13]. Identifying neurophysiological markers that reflect such changes could provide a valuable framework for understanding and optimizing tES interventions. Auditory steady-state response (ASSR) is a continuous, synchronized activity entrained by a rhythmic auditory stimulus and characterized by stability in amplitude and phase. Primarily, it reflects activity in the auditory cortex and the related neural pathways [14], [15]. Although generated within the auditory system, ASSR spreads beyond the auditory cortex [16], [17] and can be detected with the electroencephalogram (EEG) over a wide scalp area predominantly in the frontocentral region [18]. It has been shown that auditory steady-state stimulation at around 40 Hz produces the strongest and most stable ASSR [19]. As a result, 40-Hz ASSR is considered a robust way to evaluate the generation and maintenance of oscillatory gamma activity. Moreover, it is proposed as a neurophysiological marker of excitatory-inhibitory (E/I) balance [20], [21]. Cortical E/I balance represents the ratio between the activity of excitatory and inhibitory cells within neural circuits, which contributes to the integrity and excitability of the system [22], [23]. Evidence suggests that neuropsychiatric disorders characterized by aberrant E/I balance, such as schizophrenia [24], depression or autism spectrum disorder [23], also exhibit deficits in gamma ASSR [20], [26], [27], [28], making it a robust and physiologically relevant biomarker sensitive to excitability changes.

Considering that both tES action and the generation of ASSR may involve overlapping neurophysiological mechanisms, ASSR modulation via tES could provide significant insights into the network-level outcomes of tES that have been systematically overlooked. Previous studies assessed the tES effects on gamma ASSR (for review, see [29]). Generally, the application of tDCS [30], [31], [32], [33], [34] and tACS delivered at theta [35], [36], alpha [34], [36], [37], [38], [39] or gamma [33], [35], [39], [40], [41], [42] frequencies have yielded mixed results, with a number of studies also reporting no significant changes in gamma ASSR after tES. It is likely that differences in experimental designs [29] and substantial individual variability [12] contributed to inconsistent findings in previous studies. In addition, otDCS is a relatively novel tES modality, thus, no attempts have been made to explore its effects on ASSR. For these reasons, more research is needed to clarify the influence of various extrinsic and intrinsic factors on the neuromodulatory effects of tES on ASSR.

To address these gaps, the present study systematically investigates the effects of tDCS, tACS, and otDCS applied over the left posterior parietal cortex on 40-Hz ASSR in healthy adults. The main aim of the study was to explore whether ASSR could reflect tES-induced network-level changes and compare the effects across different tES modalities. We have selected tES protocols commonly applied to modulate cognitive processes such as memory and attention [43], [44], i.e., processes that have also been related to ASSRs [45]. The intensity of stimulation was selected to be comparable across tES modalities, and the electrode montage was kept consistent, inducing widely distributed electrical field. Oscillatory protocols (tACS and otDCS) were delivered at theta frequency (4-8 Hz), expecting an indirect effect on gamma synchronization reflected by ASSR due to theta-gamma coupling [35], [46].

The exploratory aim of the study was to address the potential sources of variability of tES effects. High inter-individual variability in tES outcomes can be linked to baseline neural excitability and subject-specific E/I balance [12], [47]. Thus, we explored inter-individual variability in ASSR, the variability in tES-induced ASSR changes, in relation to changes in cognitive performance under different tES conditions.

## II. Methods

### A. Study design

The data were collected from a sham-controlled, randomized cross-over experiment conducted as part of the larger MEMORYST study (for a detailed description, see [48]). Each participant completed four experimental sessions, spaced at least one week apart. In a counterbalanced order, participants received one of four stimulation conditions: tDCS, otDCS, tACS, or sham. The stimulation was administered while participants performed cognitive tasks. Immediately following tES, 5 min resting-state EEG activity was acquired, after which click-based periodic auditory stimulation was delivered during an EEG recording.

### B. Participants

A group of healthy young adults participated in the study, and all satisfied the inclusion criteria for tES application [48].

To determine the sample size, a priori power analysis for planned statistical analysis in a within-subject design was conducted for the theoretically defined medium effect size i.e. Cohen |d|= 0.50 [49]. The analysis showed that 42 participants were sufficient to detect |d| ≥ 0.50 with a power of 0.90. The participants were enrolled on a rolling basis until the desired sample size was met. The CONSORT flow chart provides a complete overview of participants’ inclusion/exclusion (Fig. 1).

**Fig. 1.**
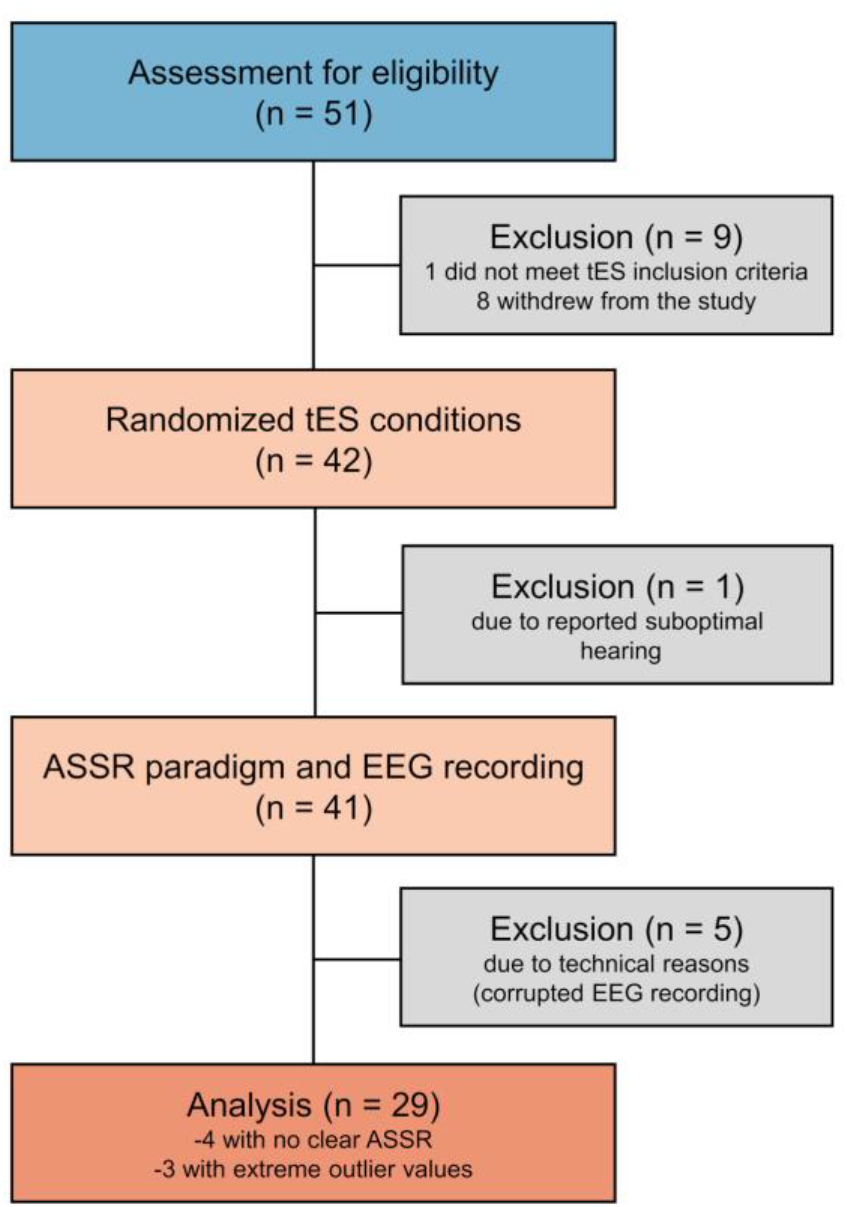
CONSORT flow chart of participant exclusion.

After the initial eligibility assessment, 42 participants completed all four tES sessions, and the ASSR paradigm was administered to 41; data of 5 participants were not analyzed due to technical reasons (missing, corrupted or interrupted EEG recording), resulting in data of 36 participants used in the analysis. Additional 7 participants were excluded during the analysis (4 showing no clear ASSR and 3 with outlier values), resulting in the final sample of 29 participants (M_age_ = 25.13, SD_age_ = 3.93; 17 female) used for statistical evaluation. Due to significant reduction in sample size, we conducted sensitivity analysis which showed the achieved power to detect |d| = 0.50 was 0.75, |d| = 0.54 will be detected with the power of 0.80, and |d| ≥ 0.62 with the power of 0.90, indicating that the statistical tests have sufficient sensitivity to detect medium-to-large effects.

The study was conducted in line with the guidelines of the Declaration of Helsinki and was approved by the Institutional Ethics Board (EO129/2020). Written informed consent was obtained from all participants and all were compensated for their participation.

### C. Transcranial Electrical Stimulation

A mobile, battery-operated, hybrid tDCS-EEG Starstim device (Neuroelectrics Inc., Barcelona, Spain) was used for both EEG acquisition and tES. The device was attached to the back of the neoprene cap and was remotely operated via Neuroelectrics® Instrument Controller (NIC) software (Neuroelectrics Inc., Barcelona, Spain).

Two round rubber electrodes (25 cm^2^) inserted in saline-soaked sponge pockets were used for stimulation. Stimulation electrodes were placed at P3 and the contralateral cheek (return). In each stimulation condition, the protocol lasted 20 minutes in total. tACS was delivered as 2 mA peak-to-peak (i.e., between −1 mA and +1 mA) sinusoidal oscillating current at individual theta frequency (ITF, 4–8 Hz). The method used for ITF extraction is described in detail in [50]. The otDCS was delivered in positive polarity (i.e., anodal current at P3) with the current oscillating at ±0.5 mA in sinusoidal mode around +1.5 mA (i.e., between +1 mA and +2 mA), also at ITF. Anodal tDCS had a constant current intensity of +1.5 mA. All active stimulation protocols had a gradual ramp up/down of 30 s at the beginning and at the end of the stimulation period. Finally, in the sham condition, the current was briefly delivered for 60 s at the beginning and at the end of the stimulation period (double ramp up/down sham protocol), both with a gradual ramp up to 1.5 mA (30 s) and an immediate gradual ramp down (30 s).

We used current flow modeling to investigate the spatial distribution and estimate the intensity of the electric field induced by tES currents in relevant cortical areas. The e-field model was calculated in SIMNIBS 4.5.0 [51], [52] on an average head model (m2m_MNI152) for current intensity of 1.5mA. The regions of interest were defined using MNI coordinates for the left PPC (−47, −71, 34), and left superior temporal gyrus (−60, −44, 6), both with a sphere radius of 10 mm. The e-field strength was 0.27 in the left PPC and 0.24 in the left superior temporal gyrus, indicating substantial field distribution in parietal and auditory cortices (Fig. 2).

**Fig. 2.**
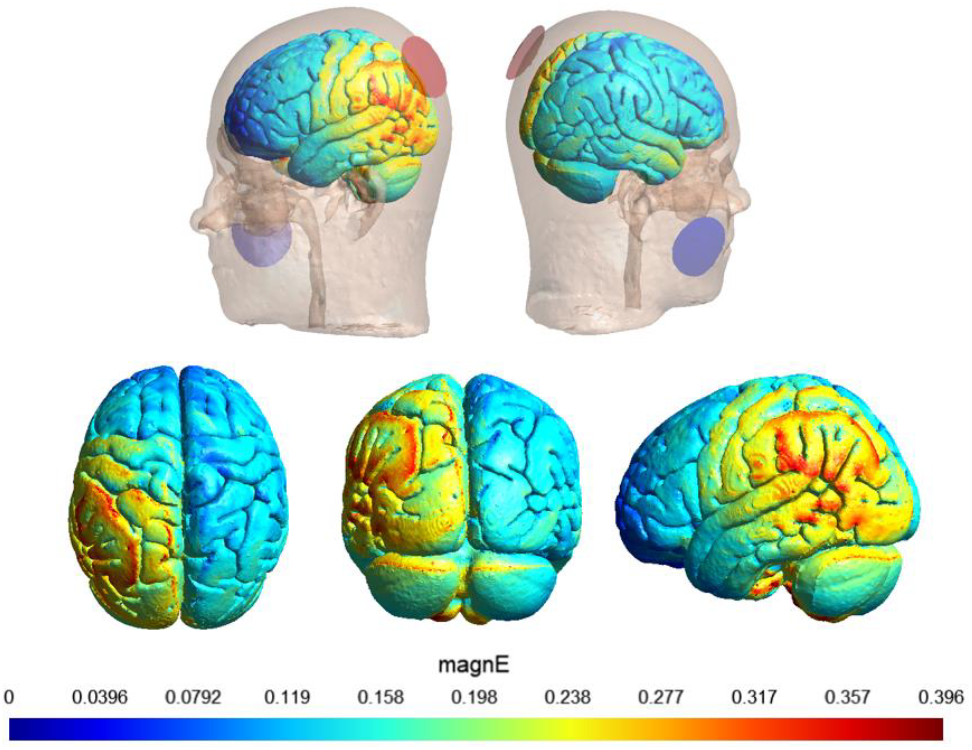
Electric field (e-field) distribution simulated using SIMNIBS 4.5.0 on the average head model for a montage with round electrodes placed over P3-contraletral cheek and intensity of 1.5 mA.

During tES, participants performed computerized cognitive tasks to keep them engaged and create uniform conditions for each participant across all tES conditions [53]. They completed a short-term associative memory (AM) task, which consisted of 42 pre-randomized sequences of 3 to 5 single digits (0–9), each paired with a colored card (red, blue, green, yellow, gray, or pink). Digits appeared sequentially in white ink for 1250 ms, with a 250 ms interval between stimuli. Participants had to memorize each digit-color pair and, when cued with one color card, recall the corresponding digit. The outcome measure was the proportion of correctly recalled numbers.

### D. EEG Acquisition

The EEG signals were recorded with a sampling rate of 500 Hz, 0–125 Hz (DC coupled) bandwidth, and 24 bits—0.05 µV resolution. Gel Ag/AgCl electrodes (4 mm diameter, 1 cm2 gel-contact area) were placed according to the international 10-10 EEG system at Fp1, Fp2, Fz, F3, F4, F7, F8, T7, T8, Cz, C3, C4, CP5, CP6, Pz, P4, PO7, PO8 and Oz positions. The 19-electrode set-up was selected to avoid contact with the stimulating electrode (P3) while maintaining reasonable whole-head coverage. For ground (DRL) and reference (CMS), either the dual CMS-DRL ear-clip electrode attached to the right earlobe, or two pre-gelled sticktrodes - the CMS on the right mastoid, and the DRL just below - were used. The manufacturer-recommended ear-clip configuration was initially assessed for signal quality in each participant. However, when due to anatomical constraints (e.g., small/narrow or pierced earlobe), this was not possible, the stickrodes were used (n = 16)

### E. Auditory Stimulation

40-Hz click-based auditory stimulus was designed in the MATLAB environment and presented binaurally through earphones with a sound pressure level adjusted to 60 dB. Each stimulus lasted 500 ms and consisted of 20 white noise clicks of 1.5 ms duration which were separated by silent periods of 25 ms, corresponding to 40 Hz. The stimulus was presented 100 times with an inter-stimulus interval set at 750 ms, resulting in an overall duration of approximately 2 minutes. Subjects were instructed to sit comfortably and listen to the sounds while looking at a fixation point on the screen in front of them.

### F. EEG Data Processing

Data preprocessing was performed in the Matlab environment using the EEGlab package [54]. Data were filtered by applying a 1-Hz highpass filter and notch filter (48-52 Hz) to remove line noise. Bad channels were manually rejected and reconstructed using spline interpolation after which channels were re-referenced to average. Independent component analysis (ICA) was performed using default EEGlab settings. Components related to eye blinks were removed. Next, data was epoched at −500 ms to 1100 ms relative to the stimulus onset, and epochs containing substantial noise or artifacts were manually removed.

Fieldtrip package [55] was used for data analysis. Morlet wavelet (10 cycles) time-frequency decomposition was performed in the frequency range of 30-50 Hz in 1 Hz steps in the time window from −500 ms to 1100 ms. Phase information was extracted to calculate the Phase Locking Index (PLI, phase consistency over epochs). To compute evoked amplitude (EA, amplitude of the evoked activity), the time series of the signals were averaged over epochs and Morlet wavelet decomposition with the same parameters was performed on the averaged signal, obtaining amplitude information. Relative baseline normalization was applied to PLI and EA data by dividing the values at each time point by the average in the baseline period (−400 to 0 ms). PLI and EA values were averaged in the time window between 200 ms and 500 ms post-stimulus onset, in the frequency range of 40±2 Hz and over channels in three frontocentral regions of interest (ROIs) separately – left (F3, C3), midline (Fz, Cz) and right (F4, C4).

### G. Statistical Evaluation

Statistical analysis was carried out using R programming language. Prior to the statistical evaluation, a visual inspection of time-frequency plots was performed to ensure the inclusion of only those participants who showed clear ASSR in the sham condition. Extreme outliers were identified using boxplots and participant-data with values above Q3 + 3×IQR or below Q1 – 3×IQR in at least one of the measures were excluded from further analysis.

To examine the effects of stimulation condition and ROI on PLI and EA values, separate linear mixed-effects models were fitted for both measures using the nlme package [56], [57] This approach was selected to account for potential individual differences often discussed in tES research [12], [47], [58] as it includes both fixed (systematic) and random (variable) effects, making it superior to repeated measures ANOVA in the presence of individual variability [59]. In addition, mixed-effects modeling helps to avoid the issue of mathematical coupling often encountered in noninvasive brain stimulation studies [60]. The fixed effects in the model included stimulation condition (sham, tDCS, otDCS, tACS), ROI (left, midline, right), and their interaction. Both variables were treated as categorical predictors, with the sham condition and midline region coded as reference levels. Random effects were specified to include both random intercepts and random slopes for stimulation condition nested within subject, allowing individual variation in both baseline PLI/EA and stimulation effects. The model was estimated using restricted maximum likelihood.

Furthermore, we computed the Pearson correlation between the random intercepts and slopes extracted from the linear mixed-effects model. To assess the significance of this correlation, we performed a non-parametric bootstrap procedure with 10,000 iterations. In each iteration, subjects were resampled with replacement, and the Pearson correlation was recalculated to generate a bootstrap distribution of correlations. A 95% confidence interval was obtained from the 2.5th and 97.5th percentiles of this distribution. The observed correlation was then compared to the bootstrap distribution to calculate a two-sided p-value.

To estimate the relationship between changes in PLI and EA induced by three different stimulation types, we computed Spearman’s correlations between simple difference scores (tDCS – sham, otDCS – sham, tACS – sham). Finally, to estimate relationship between ASSR measures and AM task outcomes, tES-induced changes in AM performance in each tES condition relative to sham (tES – sham) were related to corresponding PLI/EA simple differences (tES – sham).

## III. Results

Grand average time-frequency plots and topographies depicting baseline-corrected PLI and EA are presented in Fig. 3. A clear increase in PLI and EA is present during the auditory stimulation at the stimulation frequency (40 Hz) and the response is localized in the frontocentral area. Comparisons between conditions were performed in the left (F3, C3), midline (Fz, Cz), and right (F4, C4) ROIs separately, after averaging responses in two corresponding channels. PLI and EA simple differences between each tES condition and sham condition (tES – sham) were computed. On average, the change in PLI after each tES condition in the whole frontocentral area was close to 0 (tDCS, M = −0.03, SD = 0.56; otDCS, M = −0.01, SD = 0.57; tACS, M = −0.23, SD = 0.57). Similar results were present in EA simple difference (tDCS, M = −0.06, SD = 0.73; otDCS, M = −0.01, SD = 0.82; tACS, M = −0.29, SD = 0.79). However, a closer look revealed high inter-individual variability in effects with pronounced changes in both directions, which resulted in a net near zero change at the group level. Namely, looking at the individual level of simple differences between tES conditions, we observed 15/29 participants showing an increased PLI in the tDCS condition, 14/29 in the otDCS condition, and 14/29 in the tACS condition. PLI simple difference values among subjects ranged from −1.23 to 0.89 for tDCS, from −1.31 to 0.84 for otDCS, and from −1.46 to 0.57 for tACS. Examples of subjects that showed decreased (S1) and increased (S2) PLI relative to the sham condition following tES and grand average simple differences are shown in Fig. 4. A similar pattern was observed for EA, with 13/29 subjects exhibiting increased EA in tDCS condition, 17/29 in otDCS, and 14/29 in tACS. Individual EA simple differences ranged from −1.75 to 1.52 for tDCS, from −1.79 to 1.39 for otDCS, and from −2.04 to 1.07 for tACS.

**Fig. 3.**
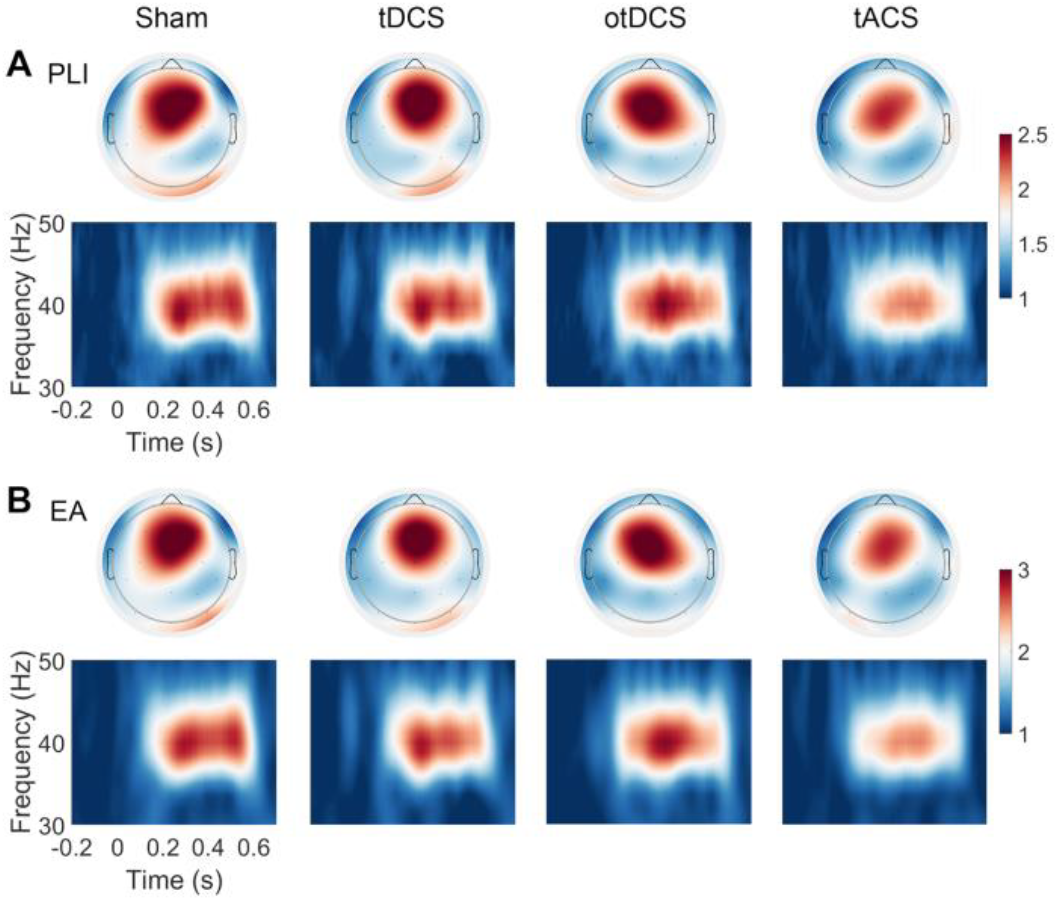
PLI (A) and EA (B) grand-average plots. Top row – topographies with average response in each channel in the time window of 200-500 ms and frequency window of 40±2 Hz; Bottom row – time-frequency plots with responses averaged over the 6 frontocentral channels (F3, Fz, F4, C3, Cz, C4) for visualization purpose. Responses in each condition (sham, tDCS, otDCS, and tACS) are presented in separate columns. Green rectangles in A depict the analyzed regions (left, midline, right).

**Fig. 4.**
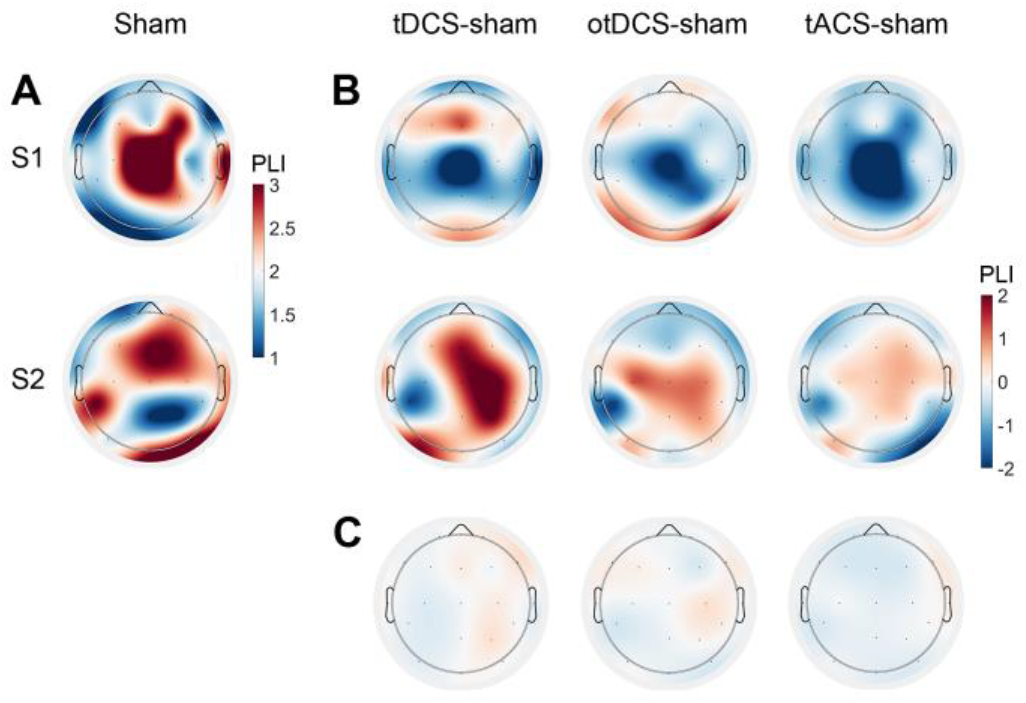
A-B. Single-subject topography plots of PLI in sham condition (A) and PLI simple difference (tES – sham) in each tES condition (B). Examples of ASSRs from participants who showed decreased PLI (S1) and increased PLI (S2) after tES are presented. C. Grand average topography plots of PLI contrast (tES – sham) in each tES condition for two subjects.

Mixed-effects linear modeling revealed consistent spatial effects across both PLI and EA measures, with significantly lower values observed in the lateral (left and right) ROIs compared to the midline ROI (PLI: β = –0.22, p < 0.01; EA: β = –0.31, p < 0.01). Stimulation condition did not have a significant main effect on either measure (PLI: β = –0.075, p = 0.12; EA: β = –0.09, p = 0.17), nor was there any significant condition-by-region interaction (PLI: β = 0.01, p = 0.77; EA: β = 0.01, p = 0.82), indicating that tES did not produce uniform modulation across regions or conditions. Baseline values, corresponding to the sham condition at the midline region, were estimated at 2.45 for PLI and 2.91 for EA as indicated by the intercept estimates. Substantial between-subject variability was evident in both measures, with random intercept SDs of 0.44 (PLI) and 0.64 (EA), and random slope SDs of 0.11 (PLI) and s0.17 (EA), reflecting individual differences in baseline levels and responsiveness to stimulation. Residual within-subject variability was moderate (SD_PLI_ = 0.57; SD_EA_ = 0.74). Notably, negative correlations between subject-level random intercepts and slopes (PLI, r = –0.52, 95% CI [−0.79 −0.19], p = 0.002); EA, r = –0.69, 95% CI [−0.86 −0.49], p < 0.001) were observed (Fig. 5), indicating that participants with lower baseline ASSR showed increased synchronization following tES.

**Fig. 5.**
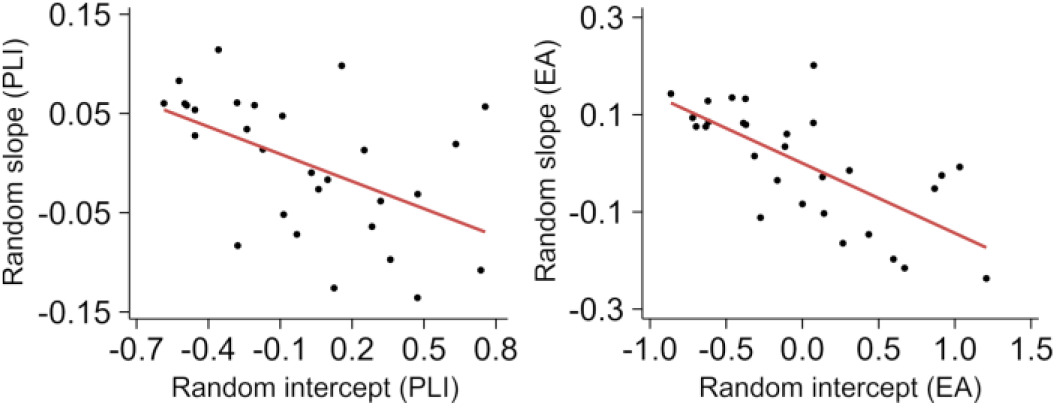
Correlation between random intercepts, representing sham ASSR level, and random slopes, representing ASSR change from sham after tES. Left, PLI (r = −0.52, p = 0.002); right, EA (r = −0.69, p < 0.001).

Significant correlations between tES-induced changes in PLI and EA relative to sham were observed between all conditions. Specifically, correlations were observed between changes induced by two oscillatory protocols (r_PLI_ = 0.38. p = 0.04; r_EA_ = 0.47, p = 0.01); as well as between tDCS-induced changes correlated with those induced by otDCS (r_PLI_ = 0.48. p = 0.01; r_EA_ = 0.38, p = 0.04) and tACS (r_PLI_ = 0.46. p = 0.01, r_EA_ = 0.50, p = 0.01). However, stimulation-to-sham changes in PLI and EA proved to be significantly and positively related to changes in AM performance only in otDCS condition (r_PLI_ = 0.46, p = 0.01; r_EA_ = 0.39, p = 0.03) (Fig. 6), while correlations in tDCS and tACS conditions did not reach statistical significance (p ≥ 0.14).

**Fig. 6.**
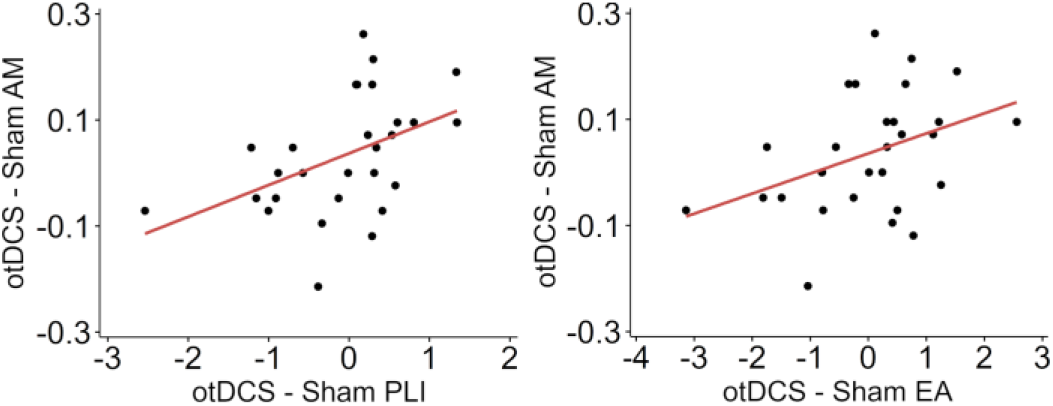
Correlations of otDCS-to-sham change in AM performance with corresponding changes in PLI (r = 0.46, p = 0.01) and EA (r = 0.39, p = 0.03).

## IV. Discussion

This study investigated the individual-specific effects of various tES protocols on 40 Hz ASSR. While group-level comparisons revealed no significant differences in ASSR between tES conditions and sham, our findings highlight considerable individual variability in response to stimulation.

Low-intensity tES techniques have shown great promise for inducing neurophysiological and behavioral changes in healthy brain [61], [62] as well as in neuropsychiatric disorders [63], [64]. However, individual variability in responsiveness continues to pose reproducibility challenges [47], [58]. This is why systematically investigating the effects of tES, especially at the neurophysiological level, is essential in order to gain a better understanding of tES mechanisms of action on the network level. To this end, extensive research is being conducted on various EEG/MEG measures to map out the neuromodulatory effects of tES [65]. The possibility of tracking changes in the standardized neurophysiological markers during or after tES is highly beneficial in this regard. One category of such measures is the steady-state responses, characterized by a stable, externally controlled oscillatory brain activity evoked by rhythmic visual, auditory, or somatosensory stimuli. In particular, ASSRs evoked by periodic auditory stimuli offer the opportunity to probe the brain’s ability to generate synchronous activity in the gamma (>30 Hz) frequency range [14], which is of particular importance due to its role in sensory and cognitive information processing [66]. In addition, ASSRs provide a unique experimental model of stable cortical electrophysiological activity [67] which is not subjected to unpredictable random variations typical for transient responses and event-related potentials [68], [69].

Building on this, our study applied different tES protocols (tDCS, otDCS, tACS) to monitor changes in ASSR magnitude and inter-trial phase synchronization, aiming to assess their potential as markers of tES neuromodulatory effects. Given the different mechanisms of action, we expected tES techniques to have differential effects on ASSR. Yet, the results revealed no significant group effect of tES on ASSR, which adds to the body of mixed evidence. Namely, previous studies reported heterogeneous tES effects on gamma-range ASSRs. With regards to tDCS, both increased [30] and reduced [31] ASSR was reported, although other studies showed no changes [32], [33], [34]. Research utilizing tACS generally showed an inhibitory effect of alpha-band tACS on ASSR [36], [37], [38], [39]; however, in one report, ASSR was enhanced following alpha-tACS in schizophrenia patients [34]. Conversely, an increase in gamma-range ASSR was shown after application of tACS at the same gamma frequency as during periodic auditory stimulation in both healthy participants [33], [40] and dyslexia patients [41], [42]. Yet, other research reported no such effects [35], [39]. So far, only two studies have investigated ASSR after theta-tACS, showing either no effects [36] or reduced ASSR [35]. Notably, we observed that ASSR in the midline region was higher compared to lateral regions, aligning with previous findings [70], [71].

The divergent findings in tES/ASSR works could be partially explained by methodological differences in tES parameters. Some studies targeted specific brain areas, including the temporal [33], [35], [36], [38], [39], [41], [42], frontal [30], [32] and central [31] areas, whereas others applied tES over multiple regions [34], [37], [40]. The low-spatial focality of tES, raises the question of neuromodulatory changes beyond the intended target area. Considering that ASSR is generated primarily in the auditory cortex [72], it might have a different sensitivity to tES depending on how far the stimulated area is from the main source of ASSR and what is the functional role of targeted areas in ASSR generation. In our study, the electrode was placed on the left PPC. As demonstrated by e-field modeling, this electrode configuration induces e-field change not only in PPC, but also in the auditory cortex, suggesting the possibility of direct modulation of ASSRs. Furthermore, PPC plays an important role in attentional control and may facilitate top-down regulation of gamma-band ASSR [73], [74]. Despite these plausible direct and indirect effects of parietal tES on ASSR, our data shows robustness of ASSR to stimulation.

In this study, we assessed the effect of individual theta frequency tES (theta-tACS and theta-otDCS) on gamma ASSR, a choice that may initially appear unconventional, although applied previously [35]. However, given the increasing interest in investigating stimulation effects beyond the primary stimulation target and ample evidence that stimulation interferes with endogenous oscillatory dynamics [4] this approach is well justified. Specifically, frequency-modulated tES has several plausible mechanisms of action. One such mechanism is cross-frequency coupling between theta and gamma oscillations [46]. Specifically, phase-amplitude coupling (PAC) between theta and gamma frequencies is a well-documented phenomenon in which the phase of theta oscillations modulates the amplitude of gamma activity, supporting cognitive functions such as working memory and attention. A recent simulation study showed that PAC can arise due to external low-frequency input [75], suggesting that theta-tACS could induce theta-gamma PAC. Although PAC presence in ASSRs itself is still debated [76], the response may be sensitive to PAC-related changes in gamma activity.

Beyond tES parameters, intrinsic factors related to brain activity per se may also contribute to inconsistent tES outcomes. Variables such as age, gender, and anatomical and physiological features of the nervous system may act as confounders, contributing to individual differences in tES outcomes [12], [47], [58]. The majority of studies that used tDCS [30], [31], [32], [33], [34] or theta-tACS [35], [36], including the present work, were characterized by relatively small samples (15-45 subjects) of healthy subjects, with the only exception of Ahn et al. [34] who recruited patients with schizophrenia. It has been suggested that tES may generally be more beneficial in psychiatric disorders as compared to healthy individuals [77]. Furthermore, tES is a relatively mild non-invasive intervention that is generally characterized by modest to moderate effect sizes [78]. In addition, healthy participants may be more resilient to external perturbations [79], which is likely applicable to ASSR due to its high individual stability [67], [80]. Accordingly, sample characteristics could have resulted in group-level null or inconsistent effects.

Moreover, large inter-individual variability exists in the cortical excitation and inhibition (E/I) balance which determines the integrity and the excitability of neural circuits [22], [23]. Accordingly, it is proposed that tES may have different effects, depending on individual-specific E/I balance [12]. For example, individuals with a more excitable state may benefit less from anodal tDCS than those with lower excitability in the stimulated area [47]. Studies also indicate that participants may be divided into groups of tES “responders” and “non-responders” due to individual features of the stimulated brain area [47], [81]. Therefore, it is possible that in the studies showing no changes in ASSR following tES, the effect could have varied between individuals, similarly to the present work.

To that note, our exploratory analyses revealed that participants showing lower baseline ASSR had an increased synchronization following tES, indicating that tES could be beneficial in neurotypical individuals with low baseline synchronization. At the same time, participants of higher baseline synchronization may benefit less, or even show lower synchronization following tES. Together, these patterns are consistent with the idea that tES may modulate neural activity in a directionally adaptive manner [82]. Given the constraints of the current study, this hypothesis should be directly tested in future research with larger samples.

The results showed that changes in ASSR induced by three tES types all correlated positively, indicating relative, although in most cases modest, consistency and overall proneness to respond to tES. This suggests that responsiveness to tES is not limited to a specific tES technique but that a responder to one type of tES is likely to show neurophysiological response on other tES techniques too. However, the magnitude of correlations between tES induced changes in ASSR indicates that there may be individuals who respond well on one, but not the other type of stimulation, and that this responsiveness may be linked to their baseline neural synchronization.

The relationship between tES-induced ASSR changes and changes in cognitive performance under different tES conditions provides additional interesting insights. Namely, we observed a significant positive relationship between tES-induced changes in neural synchronization with changes in behavioral performance; however, this relationship was observed only in otDCS condition. Specifically, otDCS-induced gains in PLI and EA were positively related to the changes in memory performance, indicating that memory improvement is potentially driven by an increase in gamma-synchronization. This finding supports the idea that otDCS may offer a synergistic mechanism, combining increased excitability of tDCS and oscillatory entrainment as in tACS [53], [83].

In sum, although no uniform effects of tES on 40-Hz ASSR were observed, the present findings emphasize the importance of individual variability. This is not surprising given that research generally shows inconsistent effects of tES on various electrophysiological parameters [84]. However, the electrophysiological and behavioral results of the present study provide preliminary indications that tES effects on 40-Hz ASSR may be individual-specific. Gamma activity arises from the interplay between glutamatergic pyramidal neurons and inhibitory GABAergic interneurons, with their balance modulating the oscillation amplitude and frequency [85]. Accordingly, ASSR may serve as a functional readout of the E/I balance [20], [21]. Given that both tES effects and ASSR generation depend on individual-specific E/I balance, the response to periodic auditory stimulation has the potential to be used as a marker of tES-related brain activity changes as well as a predictor of individual response to tES. Therefore, future studies should systematically examine the impact of physiological and behavioral individual differences on the modulatory effects of tES on gamma ASSR.

## Acknowledgment

The authors would like to thank Dunja Paunović, Katarina Vulić, Uroš Konstantinović, and Marija Stanković for their support in data collection.

